# Development of 3D breast cancer models with human T cells expressing engineered MAIT cell receptors

**DOI:** 10.1101/2022.01.29.478309

**Authors:** Madhuri Dey, Myong Hwan Kim, Momoka Nagamine, Ece Karhan, Lina Kozhaya, Mikail Dogan, Derya Unutmaz, Ibrahim T. Ozbolat

**Affiliations:** Department of Chemistry, Penn State University; University Park, PA, 16802, USA; The Huck Institutes of the Life Sciences, Penn State University; University Park, PA 16802, USA; Biomedical Engineering Department, Penn State University; University Park, PA 16802, USA; The Jackson Laboratory for Genomic Medicine; Farmington, CT 06032, USA; University of Connecticut Health Center; Farmington, CT 06032, USA; Engineering Science and Mechanics Department, Penn State University; University Park, PA 16802, USA; Materials Research Institute, Penn State University; University Park, PA 16802, USA; Department of Neurosurgery, Penn State College of Medicine, Hershey, PA 17033, USA; Penn State Cancer Institute, Penn State University, Hershey, PA 17033, USA

**Keywords:** Cancer, immunotherapy, 3D tumor models, MAIT-MR1, 3D bioprinting

## Abstract

Immunotherapy has revolutionized cancer treatment with the advent of advanced cell engineering techniques aimed at targeted therapy with reduced systemic toxicity. However, understanding the underlying immune-cancer interactions require development of advanced three-dimensional (3D) models of human tissues. In this study, we developed proof-of-concept 3D tumor models with increasing complexity to study the cytotoxic responses of CD8^+^ T cells, genetically engineered to express mucosal-associated invariant T (MAIT) cell receptors, towards MDA-MB-231 breast cancer cells. Homotypic MDA-MB-231 and heterotypic MDA-MB-231/human dermal fibroblast (HDF) tumor spheroids were primed with precursor MAIT cell ligand 5-amino-6-D-ribitylaminouracil (5-ARU). Engineered T cells effectively eliminated tumors after a 3-day culture period, demonstrating that the engineered TCR recognized MR1 expressing tumor cells in the presence of 5-ARU. Tumor cell killing efficiency of engineered T cells were also assessed by encapsulating these cells in fibrin, mimicking a tumor extracellular matrix microenvironment. Expression of proinflammatory cytokines such as IFNγ, IL-13, CCL-3 indicated immune cell activation in all tumor models, post immunotherapy. Further, in corroborating the cytotoxic activity, we found that granzymes A and B were also upregulated, in homotypic as well as heterotypic tumors. Finally, a 3D bioprinted tumor model was employed to study the effect of localization of T cells with respect to tumors. T cells bioprinted proximal to the tumor had reduced invasion index and increased cytokine secretion, which indicated a paracrine mode of immune-cancer interaction. Development of 3D tumor-T cell platforms may enable studying the complex immune-cancer interactions and engineering MAIT cells for cell-based cancer immunotherapies.

## INTRODUCTION

Cancer immunotherapy has emerged as a promising mode of treatment, harnessing the body’s immune system in eliminating cancer. In recent years, there has been remarkable progress in developing antibodies against immunological checkpoint proteins, such as CTLA-4 [1], PD-1 [2] and its ligand, PDL-1 [3]; however, success of this therapy is only limited to a subset of patients, even though these antibodies overcome the tumor-evasion mechanisms and unravel cytotoxic immune responses against tumors in certain types of cancers [4]. In addition, more recent development of cell-based immunotherapies such as Chimeric Antigen Receptor (CAR)-T cells has shown remarkable efficacy in lymphomas but had been more challenging to develop for solid tumors [5]. In both approaches, the role of tumor infiltrating T cells in generating a long-lasting anti-tumor response and the underlying immune-cancer interactions in a tumor microenvironment are poorly understood.

The tumor microenvironment consists of several cellular and acellular components including cancer cells, stromal tissue and various immune cells, such as T cells and macrophages. The composition of the microenvironment is specific to a tumor type and may also vary from patient to patient. The tumor-immune interactions in this microenvironment plays an indispensable role in cancer development, progression and control. As the dynamic tumor microenvironment is a three-dimensional (3D) structure, traditional two-dimensional (2D) cultures or animal models lack the proper tissue architecture or tumor-stroma interactions [6]. Thus, 3D cultures have emerged to be imperative tools in providing meaningful insight into disease mechanisms, and immune biomarkers, thus providing better translational relevance [7]. Some previous studies on tumor-immune interactions using 3D cultures, such as tumor spheroids, shed light on specific antitumor activity and cross-talk between natural killer cells (NK) and solid tumors. Immune-cancer co-culture in these 3D models exhibited down regulation of DNAM1 ligands on neuroblastoma cells, stimulating NK-mediated tumor killing [8], which corroborated in vivo observations. Triple negative breast cancer cell spheroids have also been used as 3D models to study the immunotherapeutic effects of a bispecific antibody against mesothelin [9].

To recognize and eliminate cancer cells, T cells must be activated by specific peptide antigens presented by major histocompatibility complex (MHC) class I molecules on the cancer cell surface. This process is difficult to recapitulate in vitro because there are very few tumorspecific antigens known to stimulate T cells, and tumor-specific T cells while critical, are rare and restricted by MHC molecules [10]. In a previous study, we developed a 2D in vitro model, in which cytotoxic mucosal-associated invariant T (MAIT) cells could be universally activated in an antigen-specific manner by employing a novel T-cell receptor engineering approach [11]. For this, a T cell subset called mucosal-associated invariant T (MAIT) cells [12], which can be activated by specific bacterial metabolites that are presented by MR1, an MHC-like molecule [13]. Because the T cell receptors (TCR) that recognize these MR1-associated metabolites are semi-invariant, T cells could be readily engineered that express synthetic versions of TCRs (Vα7.2-Jα33 associated with Vβ2) [14] using lentiviral vectors as previously described [15]. These engineered MAIT TCR (eMAIT TCR) expressing T cells are efficiently and specifically activated by cells that express MR1 and pre-cultured with bacteria supernatant that contains the riboflavin pathway metabolites such as 5-(2-oxopropylideneamino)-6-d-ribitylaminouracil (5-OP-RU) or 5-ARU [16].

In this study, we presented various 3D breast tumor models with increasing complexity to investigate the efficiency of engineered T cells in eradicating MDA-MB-231 tumors. Specifically, TCR-MR1 interaction was studied for homotypic MDA-MB-231 as well as heterotypic MDA-MB-231/HDF spheroids. eMAIT TCR expressing T cells were activated by 5-ARU ligand in the presence of MR1 expressing MDA-MB-231 spheroids or 3D models employing a hybrid approach using 3D bioprinting and also efficiently killed the MR-1 expressing tumor cells. In addition, activated T cells secreted a range of proinflammatory (IFNγ, IL-2) and anti-inflammatory (IL-6, IL-13) cytokines as well as cytotoxic mediators granzymes A and B were detected. These 3D model platforms can be a significant tool for studying the fundamental interactions between the engineered immune cells and tumors in a biomimetic 3D environment and is anticipated to evolve into an essential immune therapy model.

## MATERIALS AND METHODS

### Cells and Reagents

GFP^+^ MDA-MB-231 metastatic breast cancer cells were donated by Dr. Danny Welch, from University of Kansas, Kansas City, USA. They were cultured in RPMI 1640 media supplemented with 10% fetal bovine serum (FBS) (Life Technologies, Grand Island, NY), 1 mM Glutamine (Life Technologies, Carlsbad, CA) and 1% Penicillin-Streptomycin (Life Technologies, Carlsbad, CA). HDFs (Lonza, Walkersville, MD) were cultured in DMEM supplemented with 10% FBS, 1% glutamine, 1% sodium pyruvate, and 1% penicillin-streptomycin. HDFs were used at passages 2 through 8. PBMCs were isolated using Ficoll-paque plus (GE Healthcare, Chicago, IL). CD8^+^ T cells were purified using CD8^+^ Isolation Kit (Invitrogen, Waltham, MA). Purity (>99%) of CD8^+^ T cells were confirmed by flow cytometry staining with respective antibodies. CD8^+^ T cells were stimulated using anti-CD3/anti-CD28 coated beads (Invitrogen) at 1:2 (bead:cell) ratio and cultured in complete RPMI 1640 medium, supplemented with 10% FBS (Atlanta Biologicals, GA), and 1% Penicillin/Streptomycin (Corning Cellgro, NY) and IL-2 for 14 days. On thawing these patient derived CD8^+^ T cells, they were cultured in RPMI media supplemented with 10% FBS, 1 mM Glutamine, 1% Penicillin-Streptomycin and interleukin-2 (IL-2) at 1:500 dilution. They were cultured in suspension in 24 well plates at a density of 200,000 – 400,000 cells. Cells were all maintained at an ambient temperature of 37 °C and 5% CO_2_ flow in a humidified incubator. Cell culture medium was changed every 2-3 days. Sub-confluent cultures of adherent cells were detached from cell culture flasks using a 0.25% trypsin-0.1% EDTA solution (Life Technologies) and split to maintain cell growth.

### Engineering TCR and MR1 constructs

Design of MAIT-TCR construct was previously described [11]. Briefly, Human TCR Va7.2-Ja33 and Vβ2-Jβ2.1 gene segments were combined with mouse TCR constant alpha and β segments, respectively, and linked by a picornavirus-like 2A (p2A) self-cleaving peptide sequence. Peptide sequences of the gene segments were downloaded from Ensembl Genome Browser. Transcript ID for each gene segment and sequences for p2A and Jβ2–1 linkages were obtained from previous work [11]. The segments were connected in the following order: Vα7.2-Jα33-mTCRα-p2A-Vβ-Jβ2--mTCRβ, converted into a single open reading frame (ORF) and synthesized by Genscript. Human MR1 transcript variant 1 (NM_001531) cDNA clone was purchased from Origene (Rockville, MD). The constructs were then cloned into a lentiviral expression vector with a multiple cloning site separated from GFP reporter via an Internal Ribosomal Entry Site (IRES) [17].

### Lentiviral production and transduction

Confirmed lentiviral constructs including MR1 and eMAIT TCRs encoding vectors, were cotransfected with the packaging plasmids VSVG, pLP1 and pLP2 into HEK293T cells using Lipofectamine™ 3000 (Invitrogen) according to the manufacturer’s protocol. Lentiviral supernatants were collected, filtered and stored as previously described [15]. The cells were then transduced with multiplicity of infection of 1-5.

### Fabrication of tumor spheroids for 3D cytotoxicity studies

Two different types of tumor spheroids were used for 3D cytotoxicity studies. Homotypic tumor spheroids were comprised of ~5000 MDA-MB-231 cells, whereas, co-cultured tumor spheroids used for 3D bioprinting studies were comprised of ~5000 MDA-MB-231 cells and ~2500 HDFs. Cells were individually trypsinized and combined according to the above-mentioned ratio and seeded in 96-well U-bottom cell repellant plates (Greiner Bio-One, Monroe, NC) in 75 μl of RPMI media per well. Cells were cultured in these well plates for 2 days to form tumor spheroids.

### 3D Cytotoxicity studies on free-standing tumor spheroids

Homotypic or heterotypic tumor spheroids were treated with a MR1 ligand precursor 5-amino-6-D-ribitylaminouracil (5-ARU, Toronto Research Chemicals, Canada), diluted in RPMI media, for a period of 4 h. The compound (5-ARU) was then rinsed off from spheroids and replaced with fresh RPMI media. T cells were then introduced to spheroids at varying densities, depending on the experimental study. Briefly, for one set of experiments, the effector to target ratio (T cell to cancer cell ratio) was varied as 0.5:1, 1:1, 2:1, 4:1 and 8:1, while maintaining the concentration of 5-ARU at 3 μM. For another set of experiments, the 5-ARU concentration was varied as 30, 10, 3, 1, 0.3, 0.1, 0.03, 0.01 and 0.003 μM. The effector (T cell) to target (cancer cell) ratio, for this study, was maintained constant at 4:1. Spheroids were exposed to T cells, post 5-ARU treatment, for a period of 3 days. Images were captured daily on EVOS FL microscope (Thermo Fisher, Waltham, MA) to measure changes in fluorescence intensity of GFP^+^ tumor spheroids. To assess the viability of cancer cells after T cell treatment for 3 days, ethidium bromide (EtBr, Invitrogen) was used to stain dead nuclei. Ethidium bromide was diluted in 10 mM EDTA solution and added to spheroids and incubated for 20 min at 37 °C. Excess EtBr was carefully washed off using phosphate buffered saline (PBS), without disturbing spheroids. The spheroids were finally suspended in a small volume of ~25 μL PBS solution by mixing the cell suspension a few times using a pipette. EDTA aided in segregating spheroids into single cells. ~20 μL of this cell suspension was placed on a glass slide and covered with a cover slip to homogenously spread the solution on the glass slide. The glass slide was then mounted on the EVOS FL microscope and 15 different regions were imaged for GFP^+^ live cancer cells and EtBr stained cells. Percentage of dead cells was quantified by counting the number of EtBr stained over a sum of all nuclei using ImageJ (National Institute of Health).

### 3D Cytotoxicity studies on fibrin encapsulated cancer cells

For these studies, a poly lactic acid (PLA)-based 3D printed structure was used. This structure was designed using Autodesk Inventor (Autodesk, San Rafael, CA) and then 3D printed by Ultimaker 2 (Ultimaker, Germany). The structure had a square base and a funnel shaped mouth which was glued at the base to a coverslip, to ease microscopic visualization (Fig. S1). The square base of the 3D printed device with the raised wall ensured that the fibrin deposited in it would take up the shape of the vessel and be contained in it. The open-top funnel shape ensured that media deposited in it would also be contained for at least 3 days of culture and could be collected for further supernatant analysis. ~7500 MDA-MB-231 cells were suspended in 50 μl of 3 mg/ml fibrinogen (Sigma Aldrich, MA) and 1.2 U/ml thrombin (Sigma Aldrich). The cell suspension was then deposited inside the device and incubated at 37 °C for 30 min until complete crosslinking of fibrin. After complete crosslinking, T cells were suspended in RPMI media at an effector to target ratio of 4:1 (4xT). 10 μM 5-ARU was also added to the media. The 3D culture was maintained for a period of 3 days and then the devices were fixed for further analysis. To image the entire 3D model, Z stacks were taken on an Olympus confocal microscope (Waltham, MA). The stacks were divided equally into three zones which would represent the top, middle and bottom zones. Z stacks for each zone were about ~300 μm in thickness. The number of cells in each zone was counted using the particle analyzer function in Fiji ImageJ.

### 3D Cytotoxicity studies on fibrin encapsulated tumor spheroids

MDA-MB-231 spheroids along with engineered T cells were encapsulated in 3 mg/ml fibrinogen and 1.2 U/ml thrombin. Tumor spheroids comprised of 5000 cells/spheroid. The effector (T cell) to target (cancer cell) ratio was maintained at 4:1 (4xT). Spheroids were pre-treated with 3 μM 5-ARU for 4 h prior to fibrin encapsulation. These cultures were conducted for a period of 3 days and imaged daily on the EVOS Fl microscope to observe tumor fate after the T cell treatment. Fluorescence intensity of spheroids, before and after the T cell treatment, was then analyzed from these images using ImageJ. The invasion index of tumors was also calculated by measuring the area occupied by the T cell-treated tumors at Day 3 and dividing it by the area occupied by the non-treated control group.

### 3D Bioprinting of tumor spheroids and T cells in collagen

Heterotypic tumor spheroids comprising of MDA-MB-231 and HDFs, combined at a ratio of 2:1, was used for the 3D bioprinted tumor model. T cells were first labeled with cell tracker violet (Invitrogen) following the manufacturer’s instructions. T cell pellet, suspended in ~10 μL of culture media, was loaded into a custom-made glass pipette with a ~5μm diameter nozzle. The micropipette was prepared on a P1000 Flaming/Brown micropipette puller (Sutter Instrument, Novato, CA) using borosilicate pasteur pipettes (vWR, 14673-043, Radnor, PA). T cells were allowed to settle at the tip of the pipette ensuring a high cell density. 4 mg/ml type I collagen was used as the bioprinting substrate, which was incubated for 5 min at 37 °C, prior to T cell bioprinting. This ensured a semi-crosslinked nature of collagen that was suitable for embedded bioprinting purposes. Specifically, T cells were then bioprinted into a collagen bath in a circular pattern by continuous injection with the help of a syringe pump. After bioprinting T cells in a circular pattern, one MDA-MB-231/HDF spheroid was bioprinted at the center of the circle using aspiration-assisted bioprinting [18]. Briefly, tumor spheroids were individually aspirated from culture media using a 30G blunt nozzle, having a diameter of ~150 μm, and vacuum pressure of ~20 mmHg. They were then bioprinted into a semi-crosslinked C2F3 composite matrix at a speed of 5 mm/s. The bioprinted model was then incubated at 37 °C for another 30 min for complete crosslinking of collagen. Non-treated control groups were made by bioprinting MDA-MB-231/HDF spheroid only in the collagen bath, without T cell bioprinting. Images of T cells and MDA-MB-231/HDF spheroids were captured daily on the EVOS FL microscope using a 4x lens. Images were analyzed using Fiji ImageJ to determine changes in GFP intensity of tumor spheroids as well as invasion index of spheroids after 3 days of culture. The invasion index of tumors was calculated by measuring the area occupied by the T cell-treated tumors at Day 3 and dividing it by the area occupied by the non-treated control group.

### Cytokine analysis in supernatants

Qbeads immunoassay was used for analyzing cytokines and chemokines secreted in 3D cultures. Supernatants were collected after the end of the culture period (Day 3) and centrifuged to remove any cell debris. Supernatants were stored at −80 °C for further analysis. Capture beads fluorescently tagged with a unique signature and coated with capture antibodies directed against a specific analyte were incubated with cell culture supernatants in 96 well V-bottom plate. Once the analyte was bound by the capture beads, a fluorescent detection antibody was added to the reaction, which then bound to the analyte already bound to the beads. To maximize analyte sensitivity and reduce fluorescence background, the bead/analyte/detection was carefully washed. Data was acquired on Intellicyt iQue Screener Plus (Albuquerque, NM). The fluorescence signal was associated with the bead complex and the fluorescence intensity directly correlated to the quantity of bound analyte. Data was represented as mean fluorescence intensity. To assess the production of secreted proteins and cytokines, including granulocyte macrophage-colony stimulating factor (GM-CSF), Granzyme A, Granzyme B, IFN-α, IFN-γ, IL-1β, IL-2, IL-4, IL-6, IL-8, IL-10, IL-13, IL-17A, IL-17F, TNF-α, C-C motif chemokine Ligand 2/ monocyte chemoattractant protein 1 (CCL2/MCP1), CCL3 (MIP1a), CCL11, CX3CL1, MIG and IP10, the assay was multiplexed.

### Statistics

All data were presented as the mean ± standard deviation and analyzed by Minitab 17.3 (Minitab Inc., State College, PA) using one-way analysis of variance (ANOVA) followed by the Posthoc Tukey’s multiple comparison test. When comparing multiple groups with a single control group, a Dunnett Multiple Comparisons test was used. Statistical differences were considered significant at *p < 0.05, **p <0.01, ***p < 0.001.

## RESULTS

### Cytotoxicity induced by TCR engineered T cells on MR1 expressing free-standing homotypic MDA-MB-231 tumor spheroids

In the first tumor model (Fig. 1A), homotypic MDA-MB-231 tumor spheroids were primed with 3μM ligand precursor 5-ARU for 4 h following the T cell treatment. Human CD8+ T cells engineered to express MAIT TCRs were previously described [11]. The T cell (effector) to cancer cell (target) ratio was varied as 0.5:1 (0.5xT), 1:1 (1xT), 2:1 (2xT), 4:1 (4xT), and 8:1 (8xT). On treating the homotypic free-standing MDA-MB-231 tumor spheroids for 3 days, a dose dependent decrease in GFP intensity of tumor spheroids was observed (Fig. 1B). A ~20% decrease in GFP intensity was observed for the lowest effector density (0.5xT) as compared to a ~90% decrease for the highest effector density (8xT), as compared to the non-treated control (Fig. 1C). Additionally, ~16% cancer cell death was observed for 0.5xT, ~37% for 1xT, followed by a significant increase in cell death for 2xT (~81%) effector density onwards (Fig. 1D). Approximately, 90-95% cancer cell death was observed for 4xT and 8xT effector densities.

**Fig. 1:**
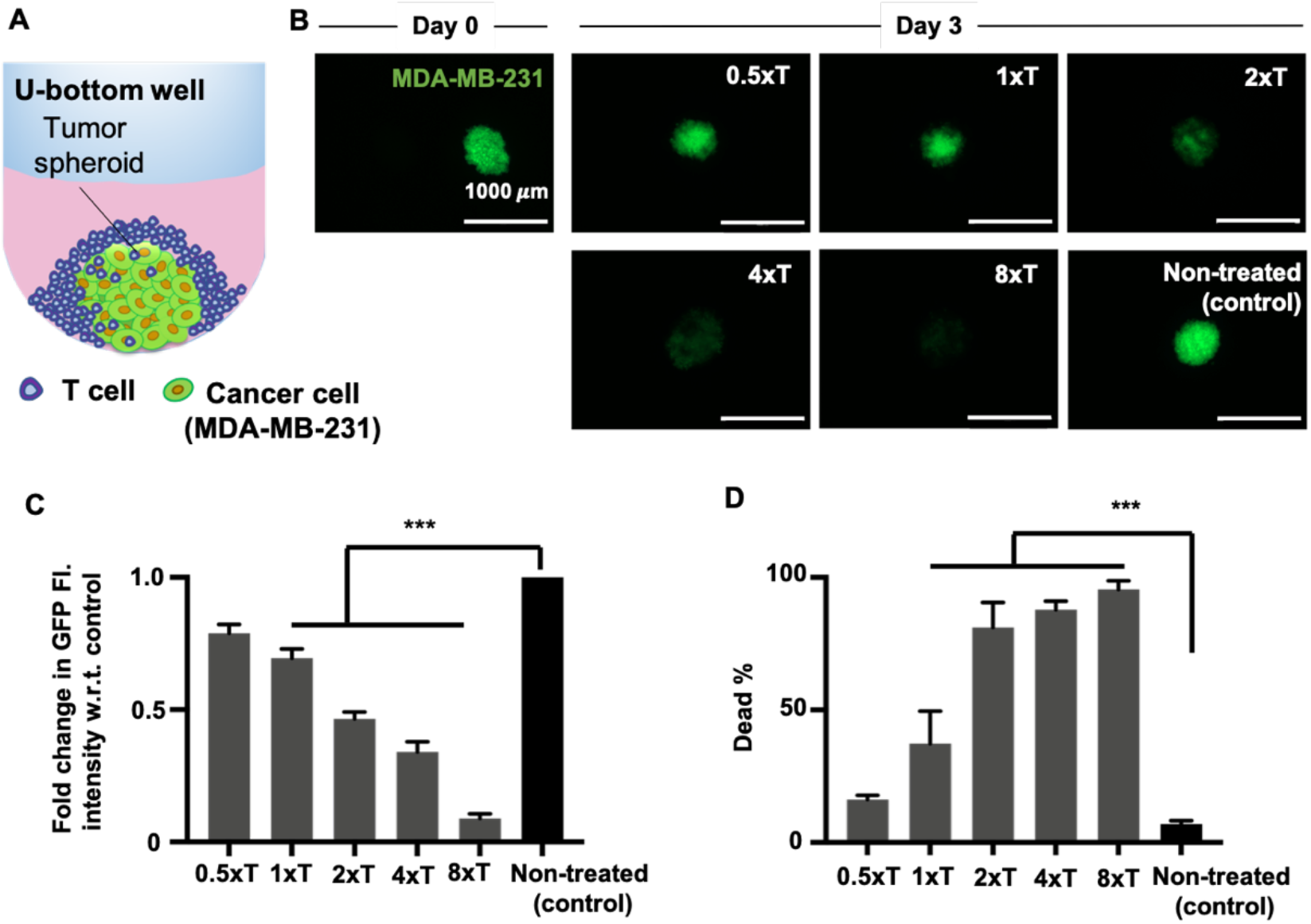
Engineered T cell mediated cytotoxicity on homotypic MDA-MB-231 tumor spheroids. (A) A schematic image of the 3D model showing a free-standing MDA-MB-231 spheroid treated with engineered T cells in a U-bottom well plate. (B) Fluorescent images of MDA-MB-231 spheroids before (Day 0) and after (Day 3) the T cell treatment. T cell (effector) to cancer cell (target) ratio was varied as 0.5:1 (0.5xT), 1:1 (1xT), 2:1 (2xT), 4:1 (4xT), 8:1 (8xT). (C) Graphical representation of fold change in GFP fluorescence intensity of MDA-MB-231 spheroids as compared to the non-treated control groups after the T cell treatment. (D) Graphical representation of MDA-MB-231 viability after the T cell treatment. (*n*=3, p*<0.05, p**<0.01, p***<0.001)

To understand the T cell-tumor crosstalk, culture supernatants collected from Day 3 of the T cell treatment were screened for the secretion of 22 cytokines and chemokines, represented on a heatmap (Fig. 2A). All T cell-treated cultures secreted granzymes A and B, as compared to the control group, with the highest secretion observed for 8xT (Fig. 2B). Similarly, there was also a dose-dependent secretion of IFNγ, CCL3, and IL-13, increasing with higher effector densities (Figs. 2C-E). Expression levels of these cytokines in the control group were extremely low or undetectable. In contrast, IL-6 and IL-8 was secreted by both treated as well as the control group. However, expression levels of IL-6 were significantly higher for the T cell-treated groups, irrespective of effector densities (Fig. 2F). Interestingly, IL-8 secretion was significantly higher for 0.5xT and 1xT effector densities and lower for 2xT, 4xT and 8xT effector densities, as compared to the control group (Fig. 2F). A dose-dependent decrease in MIG and corresponding increase in GM-CSF expression was also observed (Figs. 2G, H).

**Fig. 2:**
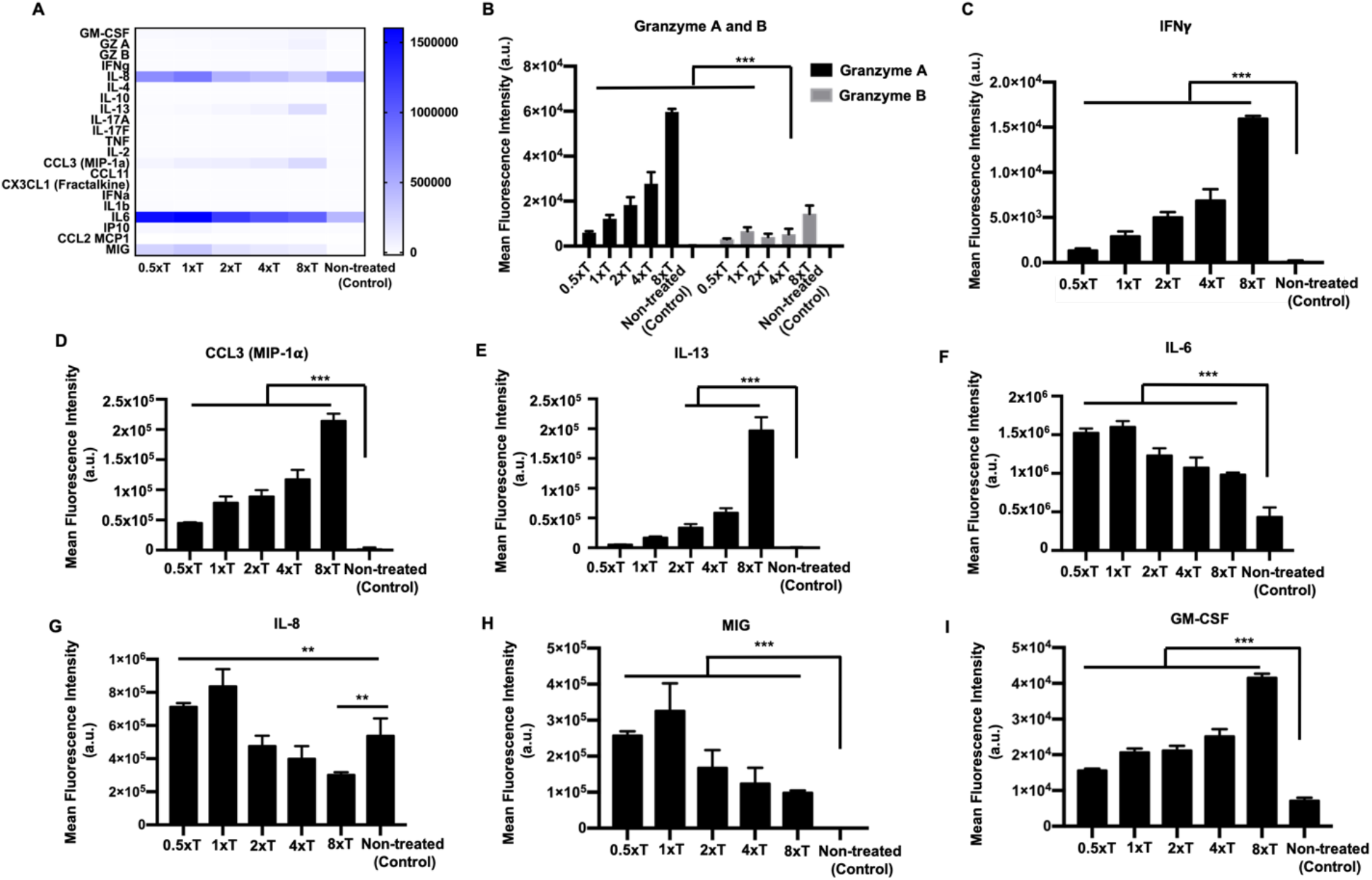
Supernatant analysis of homotypic free-standing MDA-MB-231 tumor spheroids. (A) Heatmap of all the cytokines and chemokines released in the supernatant after the T cell treatment. Graphical representation of the mean fluorescence intensity of the cytokines and chemokines secreted including (B) Granzyme A and B, (C) IFNγ, (D) CCL3, (E) IL-13, (F) IL-6, (G) IL-8, (H) MIG and (I) GM-CSF (*n*=3, p*<0.05, p**<0.01, p***<0.001).

Next, MDA-MB-231 spheroids were primed with varying 5-ARU concentrations, keeping the effector to target ratio constant at 4xT. 5-ARU concentration was varied from a low of 0.003 μM to a high of 30 μM. Control groups included spheroids treated with an inactive compound (6-FP), and non-treated spheroids. Engineered T cells exhibited a 5-ARU concentration dependent cytotoxicity on MDA-MB-231 spheroids. From 95 to 100% cancer cell death was observed for 3, 10 and 30 μM 5-ARU primed spheroids (Fig. 3A), with the lowest being ~40% death for 0.003 μM. In contrast, MDA-MB-231 spheroids exposed to 6-FP (inactive compound) and treated with 4xT effector density exhibited only ~10-15% cancer cell death, close to the non-treated control group. Immune response after the T cell treatment also followed a concentration dependent pattern. 5-ARU primed T cell-treated spheroids expressed significantly high levels of granzymes and IFNγ (Figs. 3C, D), as compared to 6-FP primed spheroids. Both, 6-FP and 5-ARU primed MDA-MB-231 spheroids exhibited secretion of IL-13, IL-6, CCL3 and GM-CSF (Figs. 3E-H). Only 3, 10 and 30 μM 5-ARU primed spheroids secreted IL-13 significantly higher than 6-FP primed spheroids. Interestingly, there was no significant difference in IL-6 secretion among all the treated and control groups. Additionally, IL-2 secretion was only observed for 30 to 0.03 μM 5-ARU primed spheroids (Fig. 3I).

**Fig. 3:**
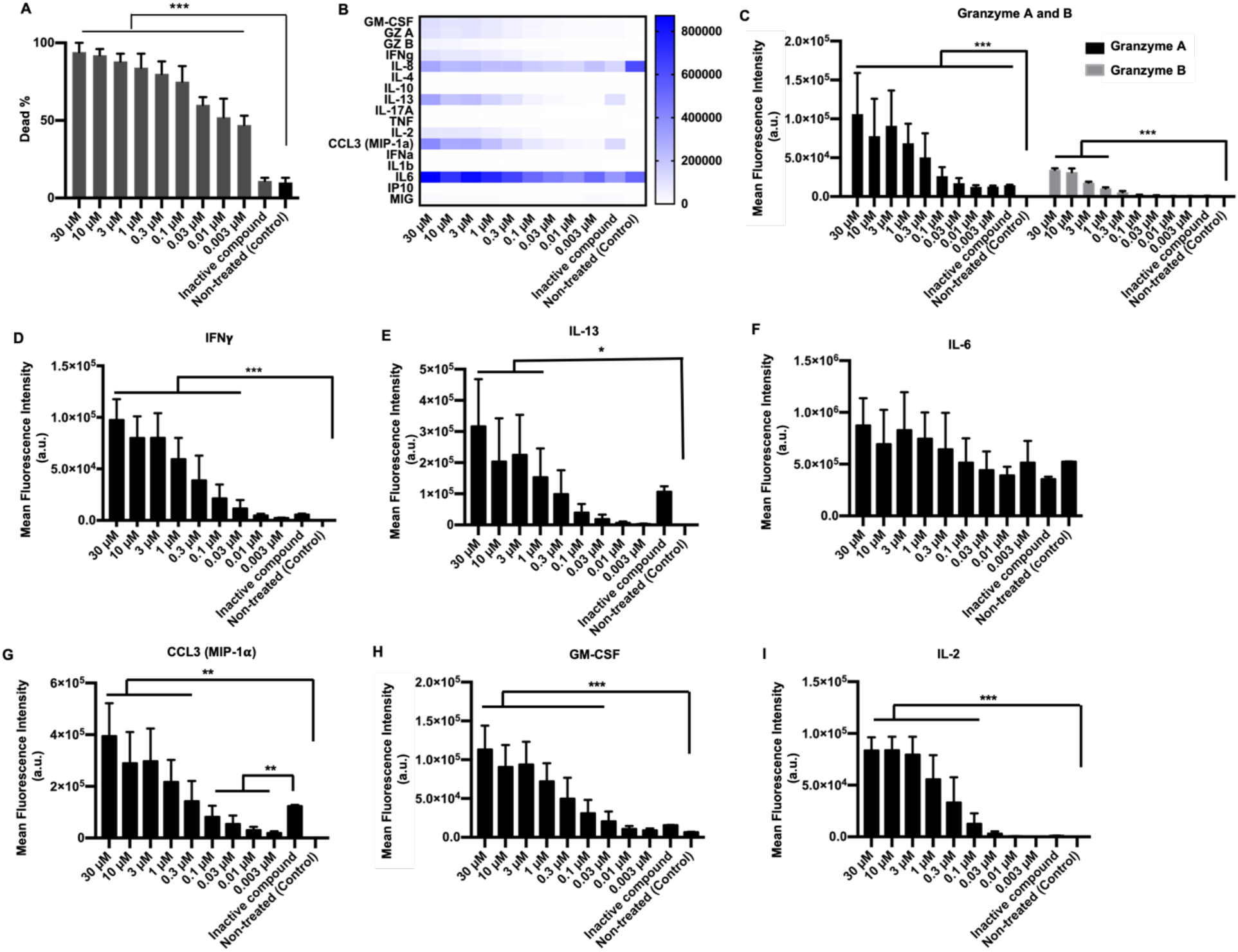
Effect of 5-ARU on engineered T cell-mediated cytotoxicity on homotypic free-standing MDA-MB-231 tumor spheroids. (A) Graphical representation of MDA-MB-231 tumor viability after the T cell treatment with varying concentrations of 5-ARU. 5-ARU concentration was varied from 30 M to 0.003 μM. (B) Heatmap of all the cytokines and chemokines released in the supernatant after the T cell treatment. Graphical representation of the mean fluorescence intensity of cytokines and chemokines secreted including (C) Granzyme A and B, (D) IFNγ, (E) IL-13, (F) IL-6, (G) CCL3, (H) GM-CSF and (I) IL-2 (*n*=3, p*<0.05, p**<0.01, p***<0.001).

### Cytotoxic response of TCR engineered T cells to MR1 expressing MDA-MB-231 cells encapsulated in fibrin

A tumor is typically surrounded by a dense matrix in its native microenvironment, comprising of extracellular matrix (ECM) proteins like fibrin or collagen. Thus, for effective anti-tumor activity and treatment response, engineered T cells need to navigate through the dense matrix to reach their target. In order to assess the efficiency of T cell killing in complex 3D microenvironments, MDA-MB-231 cells were encapsulated in fibrin and exposed to engineered T cells for a period of 3 days. Engineered T cells were deposited on top of the fibrin construct, as a second tumor model (Fig. 4A). T cells were live-imaged for a period of 16 h and observed migrating into the fibrin network (Supplementary Video S1). However, their extent of migration and penetration depth varied with varying depth of the 3D tumor model. Specifically, most T cells were seen concentrated in the top 300 μm, and their penetration depth decreased with increasing thickness of the tumor model. Due to this differential migration of T cells, the tumor model, as shown in Fig. 4A, was divided into three zones, namely top, middle and bottom zone, each zone being about ~300 μm in depth. The top zone in T cell-treated cultures showed significant reduction in GFP^+^ cells, as compared to middle or bottom zones, on Day 3 (Fig. 4B). All zones for the non-treated control group exhibited an increase in GFP intensity by Day 3. There was a significant decrease in cancer cell density for the top (~ 80%) and middle (~ 40%) zones, as compared to the control group. In contrast, the bottom zone exhibited an increase in GFP intensity owing to cancer cell proliferation (Supplementary Video S2). Cytokines and chemokines secreted after the T cell treatment were plotted on a heatmap, as shown in Fig. 5A. Specifically, granzymes, IFNγ, IL-13, CCL3, and MIG were only secreted by T cell-treated cultures (Figs. 5B-F). IL-6, IL-8, and GM-CSF were secreted by both treated and non-treated groups; however, secretion of these factors was significantly higher after the immune treatment (Figs. 5G-I).

**Fig. 4:**
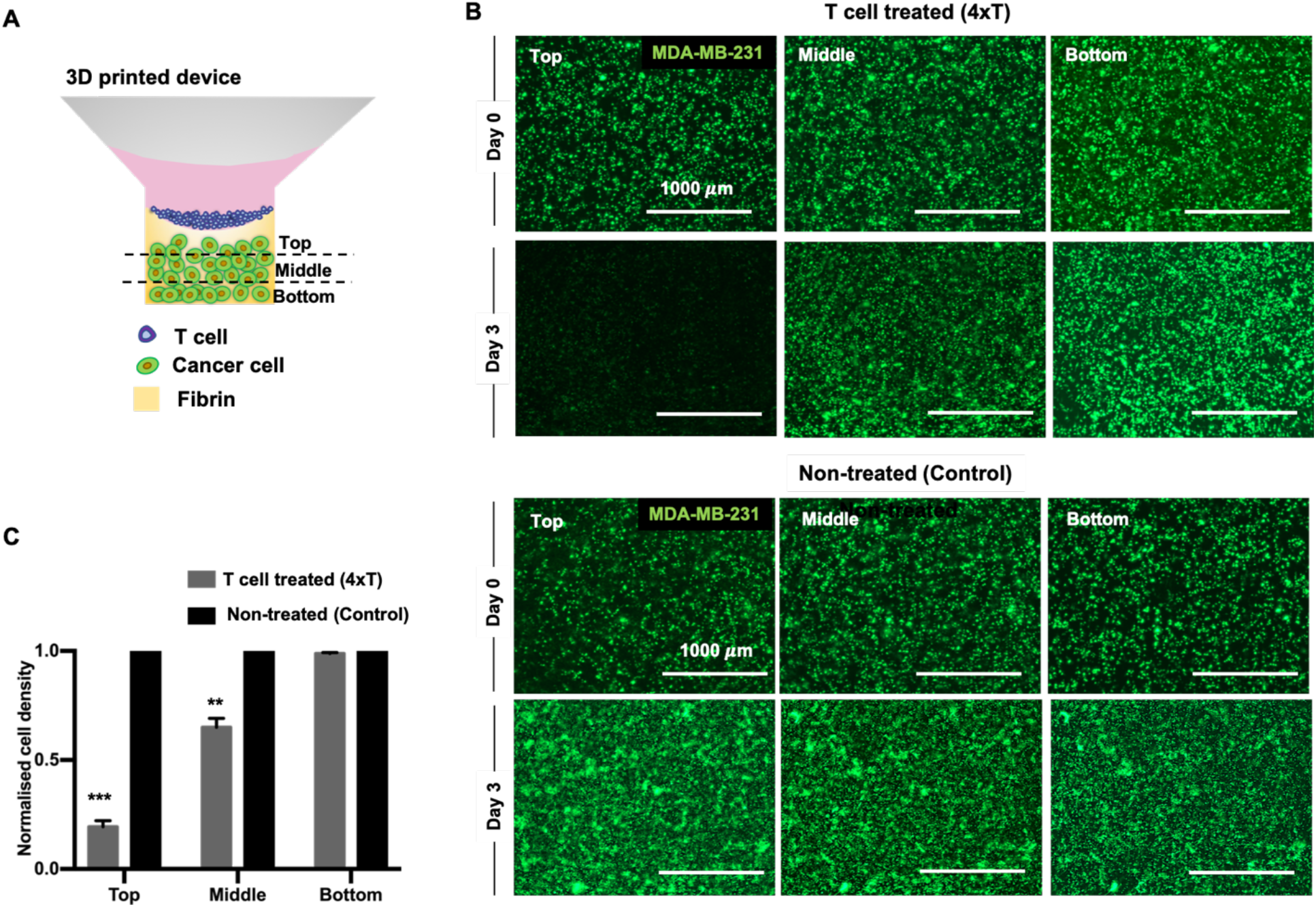
Engineered T cell mediated cytotoxicity on MDA-MB-231 cells encapsulated in fibrin. (A) A schematic image of the 3D model showing T cells deposited on top of a fibrin construct containing MDA-MB-231 cells. The 3D model was divided into top, middle and bottom zones for post T cell treatment analysis. (B) Fluorescent images of MDA-MB-231 cells before (Day 0) and after (Day 3) the T cell treatment for T cell-treated and non-treated cultures. The T cell (effector) to cancer cell (target) ratio was maintained constant at 4:1 (4xT). (C) Graphical representation of normalized MDA-MB-231 cell density in top, middle and bottom zones, after the T cell treatment. Cell densities were normalized to control samples (*n*=3, p*<0.05, p**<0.01, p***<0.001).

**Fig. 5:**
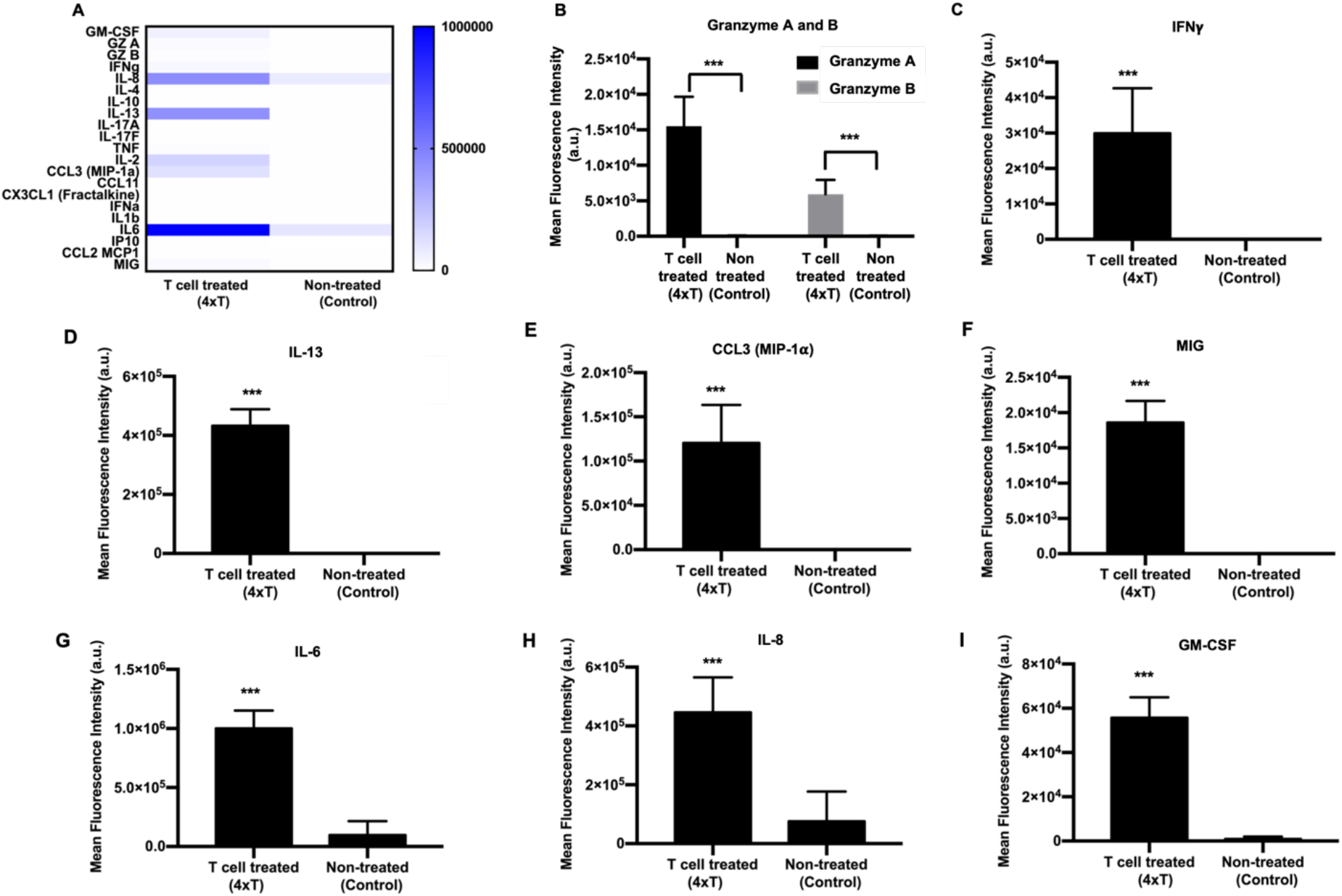
Supernatant analysis for MDA-MB-231 cells encapsulated in fibrin, post T cell treatment. (A) Heatmap of all the cytokines and chemokines released in the supernatant after the treatment. Graphical representation of the mean fluorescence intensity of cytokines and chemokines secreted including (B) Granzyme A and B, (C) IFNγ, (D) IL-13, (E) CCL3, (F) MIG, (G) IL-6, (H) IL-8 and (I) GM-CSF (*n*=3, p*<0.05, p**<0.01, p***<0.001).

### Cytotoxicity induced by TCR engineered T cells on MR1 expressing MDA-MB-231 spheroids encapsulated in fibrin

Having assessed the efficiency of engineered T cells in killing cancer cells encapsulated in fibrin, in the third tumor model, we sought to understand the effect of these cells on 3D tumor spheroids in fibrin. T cells were encapsulated in fibrin along with the MDA-MB-231 tumor spheroids to maximize their interactions (Fig. 6A). T cells were able to navigate through the fibrin network in tandem with cancer cell proliferation (Supplementary Video S3). Over 3 days of culture, T cells completely eradicated MDA-MB-231 spheroids as compared to the non-treated control group, where cancer cells proliferated from the tumor and invaded into the surrounding fibrin matrix. ~80% decrease in GFP intensity was observed after the immune treatment as compared to nontreated control group (Fig. 6C). As the cancer cells proliferated out (Supplementary Video S3), engineered T cells were able to stop their further migration and induce apoptosis on TCR-MR1 recognition. ~80% decrease in cancer invasion was observed for the T cell-treated group as compared to the control group (Fig. 6D). Secretion levels of cytokines and chemokines were depicted on a heatmap (Fig. 7A). Detectable levels of granzymes, IFNγ, CCL3, IL-13, and, MIG secretion were only observed for the T cell-treated groups (Figs. 7B-G). IL-6, IL-8 and GM-CSF were secreted by both the treated as well as non-treated groups (Figs. 7F-I). However, the secretion levels after the immune treatment were significantly higher as compared to the non-treated control group.

**Fig. 6:**
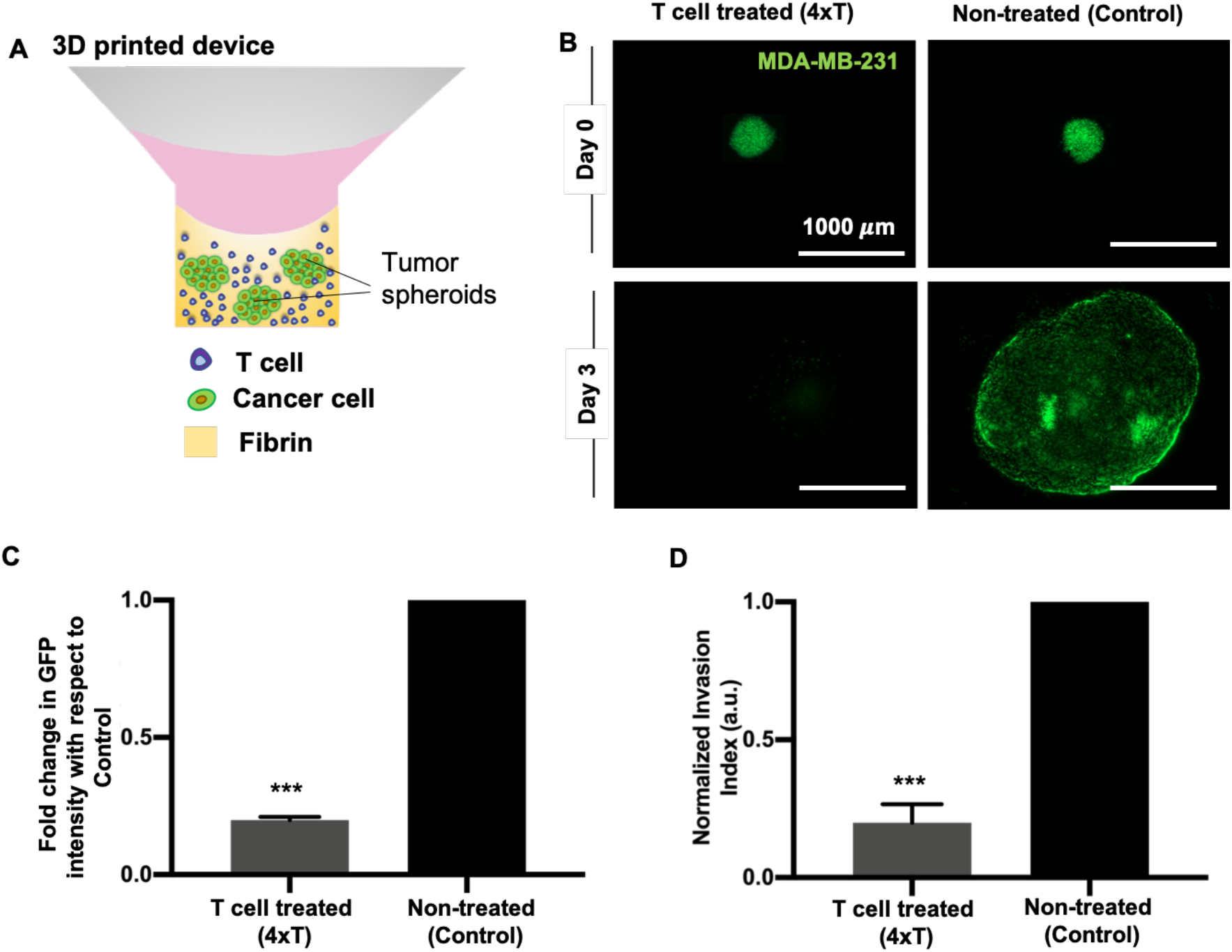
Engineered T cell-mediated cytotoxicity on homotypic MDA-MB-231 tumor spheroids encapsulated in fibrin. (A) A schematic image of the 3D model showing T cells encapsulated in fibrin along with MDA-MB-231 spheroids. (B) Fluorescent images of MDA-MB-231 spheroids before (Day 0) and after (Day 3) the T cell treatment for treated and non-treated (control) groups. The T cell (effector) to cancer cell (target) ratio was maintained constant at 4:1 (4xT). Graphical representation of (C) fold change in GFP intensity and (D) normalized invasion index of MDA-MB-231 spheroids with respect to the control, post T cell treatment (*n*=3, p*<0.05, p**<0.01, p***<0.001).

**Fig. 7:**
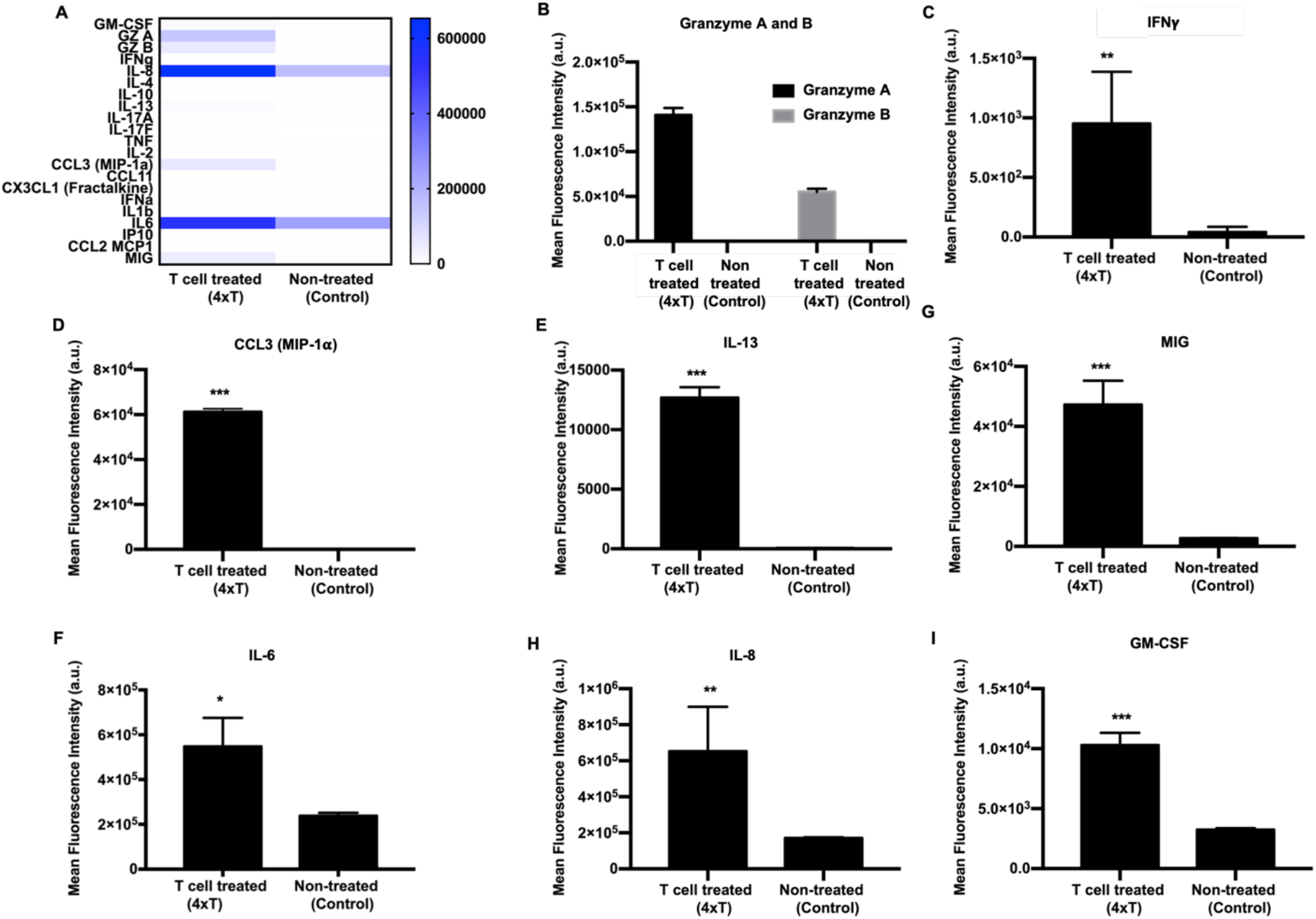
Supernatant analysis for MDA-MB-231 spheroids encapsulated in fibrin, post T cell treatment. (A) Heatmap of all cytokines and chemokines released in the supernatant after treatment. Graphical representation of the mean fluorescence intensity of cytokines and chemokines secreted including (B) Granzyme A and B, (C) IFNγ, (D) CCL3, (E) IL-13, (F) MIG, (G) IL-6, (H) IL-8 and (I) GM-CSF (*n*=3, p*<0.05, p**<0.01, p***<0.001).

### Cytotoxicity induced by TCR engineered T cells on MR1 expressing free-standing heterotypic MDA-MB-231/HDF tumor spheroids

To understand the efficiency of engineered T cell mediated cytotoxicity, heterotypic tumor spheroids comprising of MDA-MB-231 and HDFs, as the fourth tumor model, were treated with varying densities of T cells (Fig. 8A). Effector to target ratio was varied as 0.5xT, 1xT, 2xT, 4xT and 8xT (Fig. 8B). Similar to the homotypic tumor spheroids, a dose dependent increase in cytotoxicity was also observed for the heterotypic tumor spheroids, resulting in decreased GFP intensity with increasing effector densities (Fig. 8B). Significant decrease in GFP intensity was only observed for 2xT, 4xT and 8xT treatment densities, with ~60% decrease for the highest density (8xT), as compared to the non-treated control. The extent of cancer cell death in the heterotypic tumor model was lower as compared to the homotypic model. Briefly, ~20-40 % cancer cell death was observed for 0.5xT, 1xT and 2xT, ~50% for 4xT and ~70% for 8xT treatment densities (Fig. 8D). Secreted cytokines and chemokines post immune treatment were depicted on a heatmap (Fig. 9A). Production of granzymes, IFNγ, IL-13 and MIG was detectable in the T cell-treated groups, similar to all the previous tumor models (Figs. 9B-E). CCL3, IL-6, CCL2 and IL-8 was secreted by both the treated and non-treated groups (Figs. 9F-I). There was no difference in IL-6 secretion between treated and non-treated control groups. GM-CSF was also significantly produced by both treated and non-treated control groups, with a surge in secretion for the highest effector density (8xT) (Fig. S2). Interestingly, secretion of IL-8 and CCL2 were higher in the nontreated group as compared to the treated group.

**Fig. 8:**
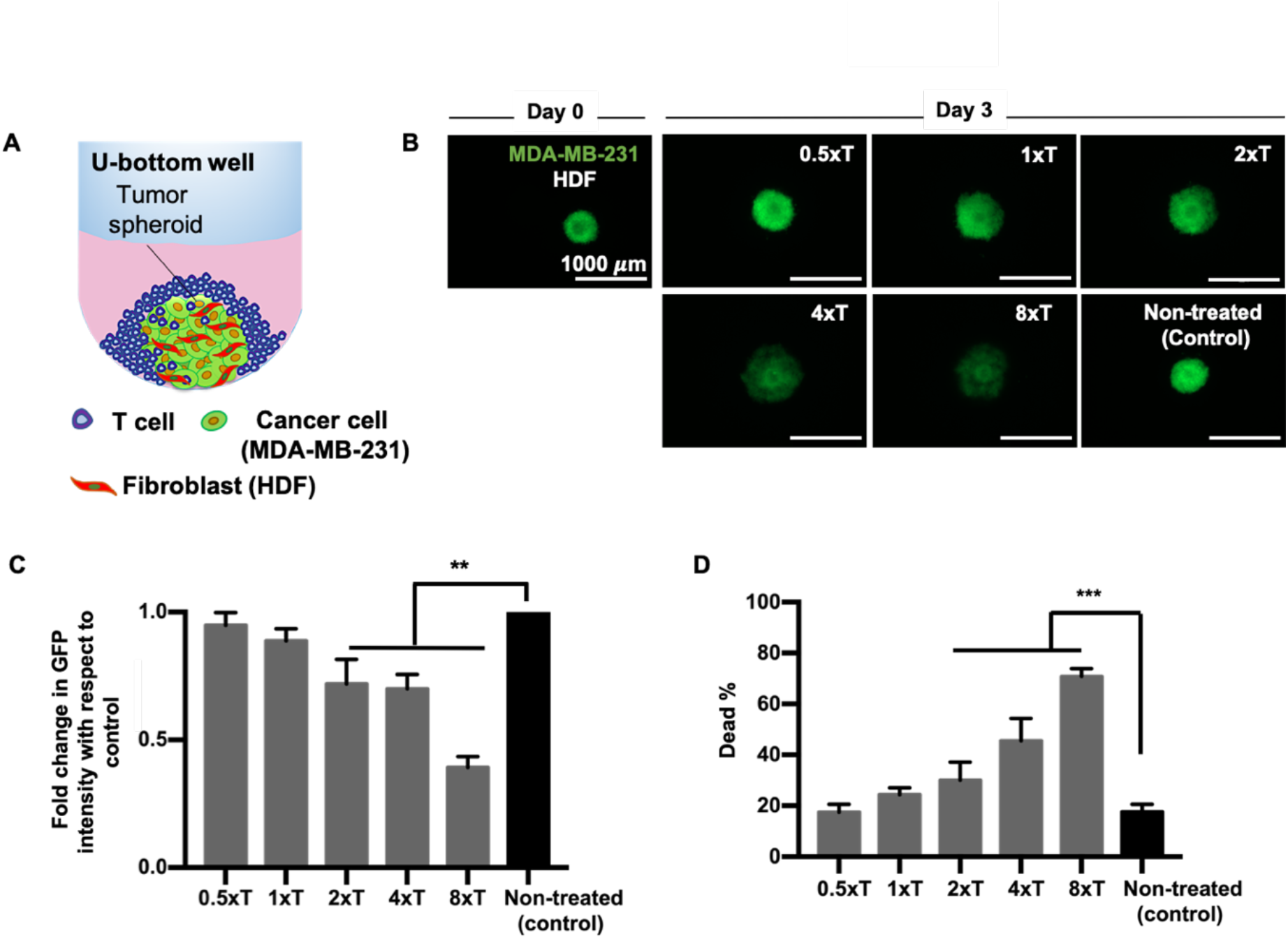
Engineered T cell mediated cytotoxicity on heterotypic free-standing MDA-MB-231/HDF tumor spheroids. (A) A schematic image of the 3D model showing free-standing MDA-MB-231/HDF spheroids treated with engineered T cells in U-bottom wells. (B) Fluorescent images of MDA-MB-231/HDF spheroids before (Day 0) and after (Day 3) the T cell treatment. T cell (effector) to cancer cell (target) ratio was varied as 0.5:1 (0.5xT), 1:1 (1xT), 2:1 (2xT), 4:1 (4xT), and 8:1 (8xT). (C) Graphical representation of fold change in GFP intensity of heterotypic spheroids as compared to non-treated control spheroids after the T cell treatment. (D) Graphical representation of tumor viability after the T cell treatment (*n*=3, p*<0.05, p**<0.01, p***<0.001).

**Fig. 9:**
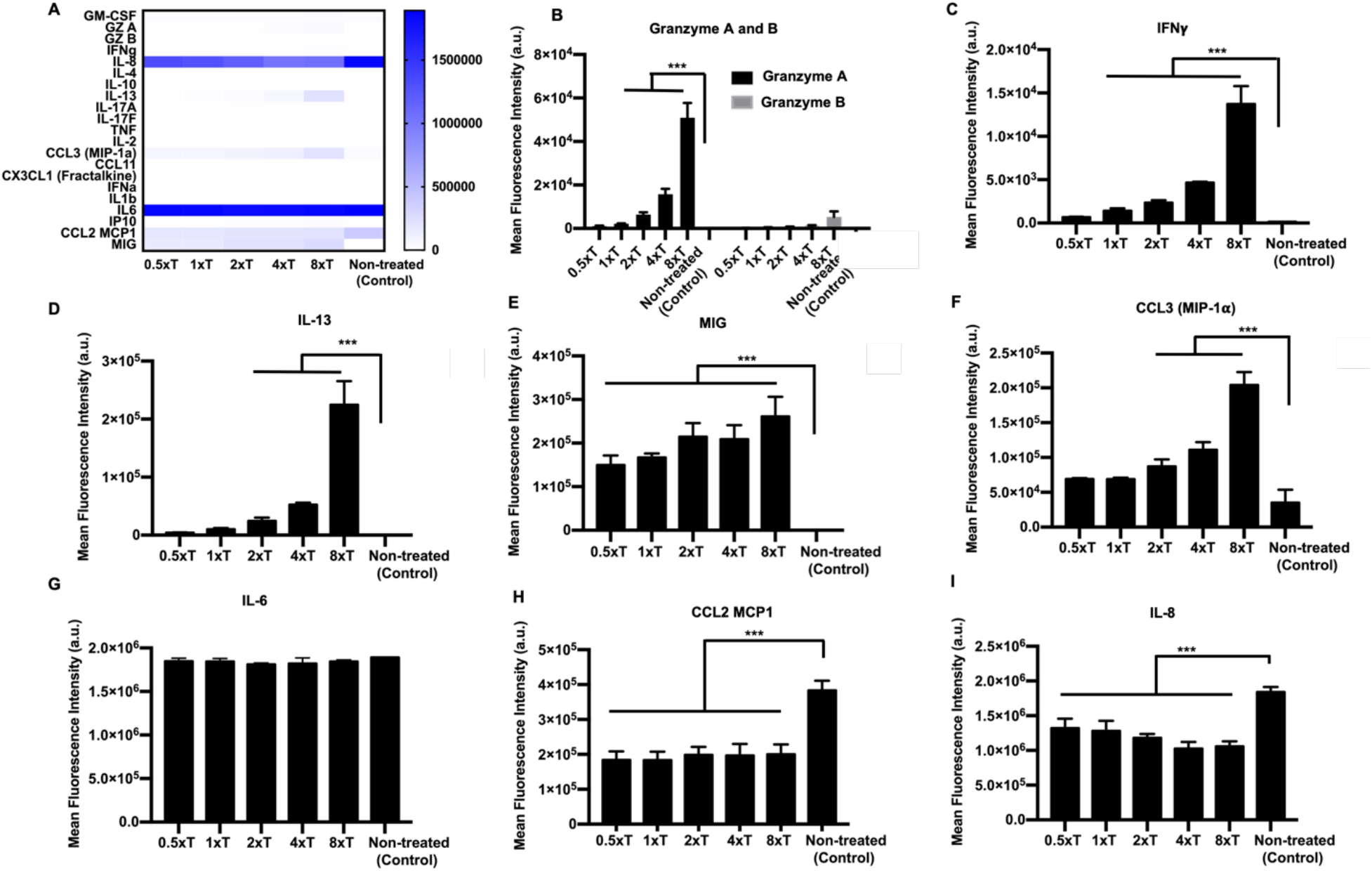
Supernatant analysis for MDA-MB-231/HDF spheroids encapsulated in fibrin, post T cell treatment. (A) Heatmap of all cytokines and chemokines released in the supernatant after the treatment. Graphical representation of the mean fluorescence intensity of cytokines and chemokines secreted including (B) Granzyme A and B, (C) IFNγ, (D) IL-13, (E) MIG, (F) CCL3, (G) IL-6, (H) CCL2 and (I) IL-8 (*n*=3, p*<0.05, p**<0.01, p***<0.001).

### Cytotoxicity induced by bioprinted TCR engineered T cells on MR1 expressing 3D bioprinted heterotypic MDA-MB-231/HDF tumor spheroids

A more complicated tumor model was also fabricated via 3D bioprinting using heterotypic MDA-MB-231/HDF tumor spheroids in order to demonstrate the effect of localization on immune-cancer interactions. T cells were loaded in a syringe and bioprinted in circular patterns into a 4 mg/ml collagen matrix. Collagen was chosen as the matrix due to its slower crosslinking at room temperature as compared to fibrin, which makes it favorable for bioprinting. After T cell bioprinting, aspiration-assisted bioprinting was employed to deposit heterotypic MDA-MB-231/HDF spheroid at the center of circular patterns. Circular patterns were developed in two ways, a smaller circle, where the T cells were located at an average distance ~250 μm from the centrally bioprinted tumor (proximal, Fig. 10A1), and a larger circle where the T cells were located at an average distance ~650 μm from the centrally bioprinted tumor (distal, Fig. 10A2). Cancer cells from the centrally bioprinted tumor spheroid proliferated and invaded into the surrounding collagen matrix for both the proximal and distal conditions (Figs. 10A1, A2). However, cancer cell invasion was ~70% lower for the proximal and ~50% lower for the distal as compared to the non-treated control group (Fig. 10B, Fig. S2). Additionally, there was a ~50-60% decrease in GFP intensity for both the proximal and distal conditions (Fig. 10C). Secreted cytokines and chemokines from treated and non-treated control groups were depicted as a heatmap (Fig. 11A). Interestingly, expression levels of granzymes, IFNγ, and CCL2 were all significantly higher for the proximal condition where T cells were bioprinted at a ~250 μm distance from the tumor periphery as compared to distal condition (Figs. 11B-D). IL-13 and MIG expression was high in both proximal and distal conditions as compared to the control group (Figs. 11E, F). Additionally, IL-6, CCL3 and GM-CSF were produced by both treated and non-treated groups, with a significantly higher expression for T cell-treated groups (Fig. 11F-I). IL-6 was equivalently expressed by all groups (Fig. 11G). The expression levels of CCL3 and GM-CSF were higher in the proximal condition as compared to distal; however, the difference was not statistically significant.

**Fig. 10:**
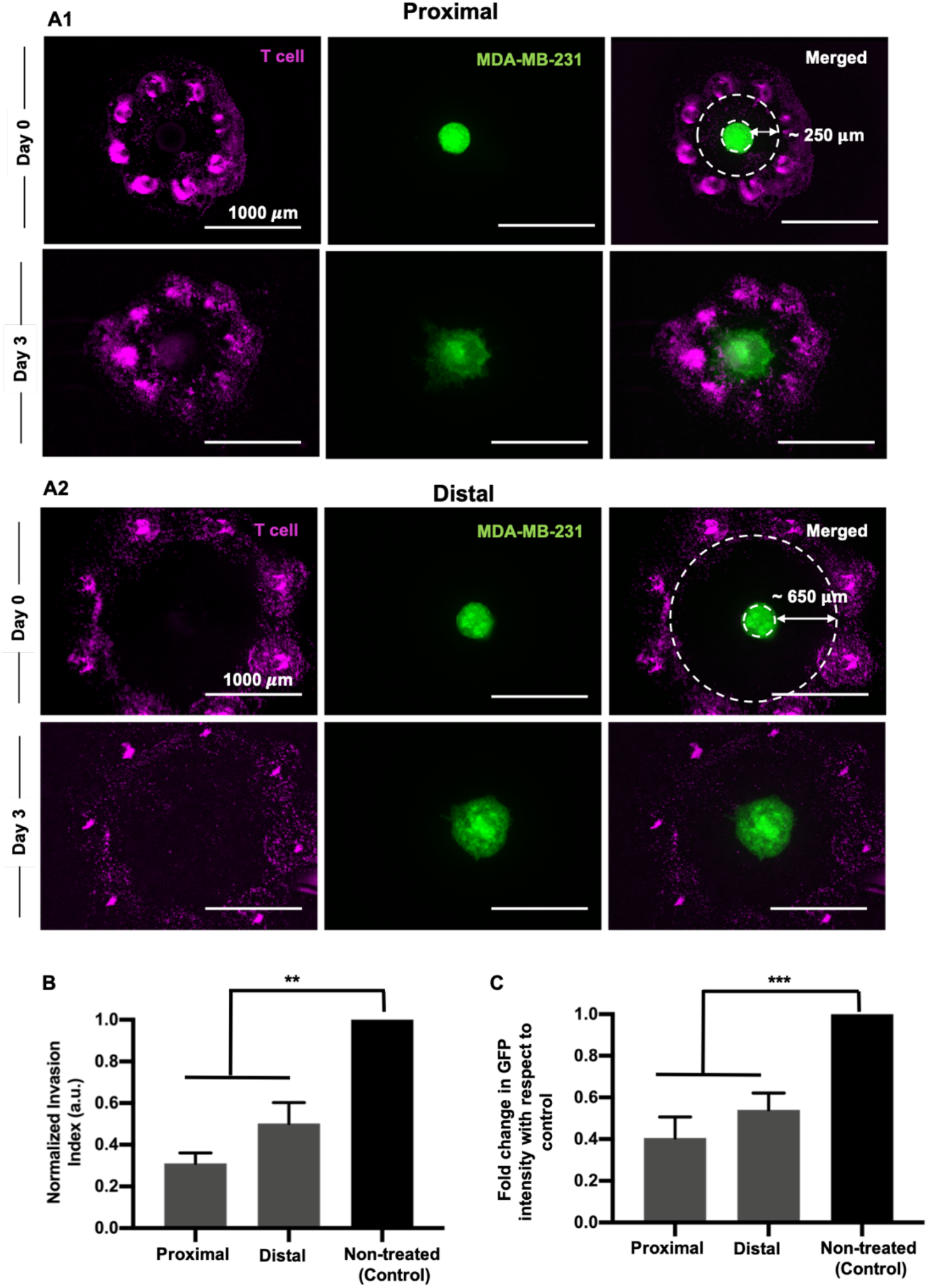
A 3D bioprinted immune-cancer model. T cells were labeled with cell tracker violet and bioprinted in a circular pattern into a collagen bath. MDA-MB-231/HDF tumor spheroids were then bioprinted at the center of the circle. The distance of the T cells from the periphery of the tumor spheroid was maintained at ~250 μm (proximal) and ~650 μm (distal). Fluorescent images of the (A1) proximally and (A2) distally bioprinted tumor models right after bioprinting (Day 0), and post T cell treatment (Day 3). (B) Graphical representation of fold change in GFP intensity of tumor spheroids bioprinted proximally or distally with respect to the T cell circle on Day 3. (D) Graphical representation of normalized invasion index of spheroids with respect to the non-treated control group on Day 3 (*n*=3, p*<0.05, p**<0.01, p***<0.001).

**Fig. 11:**
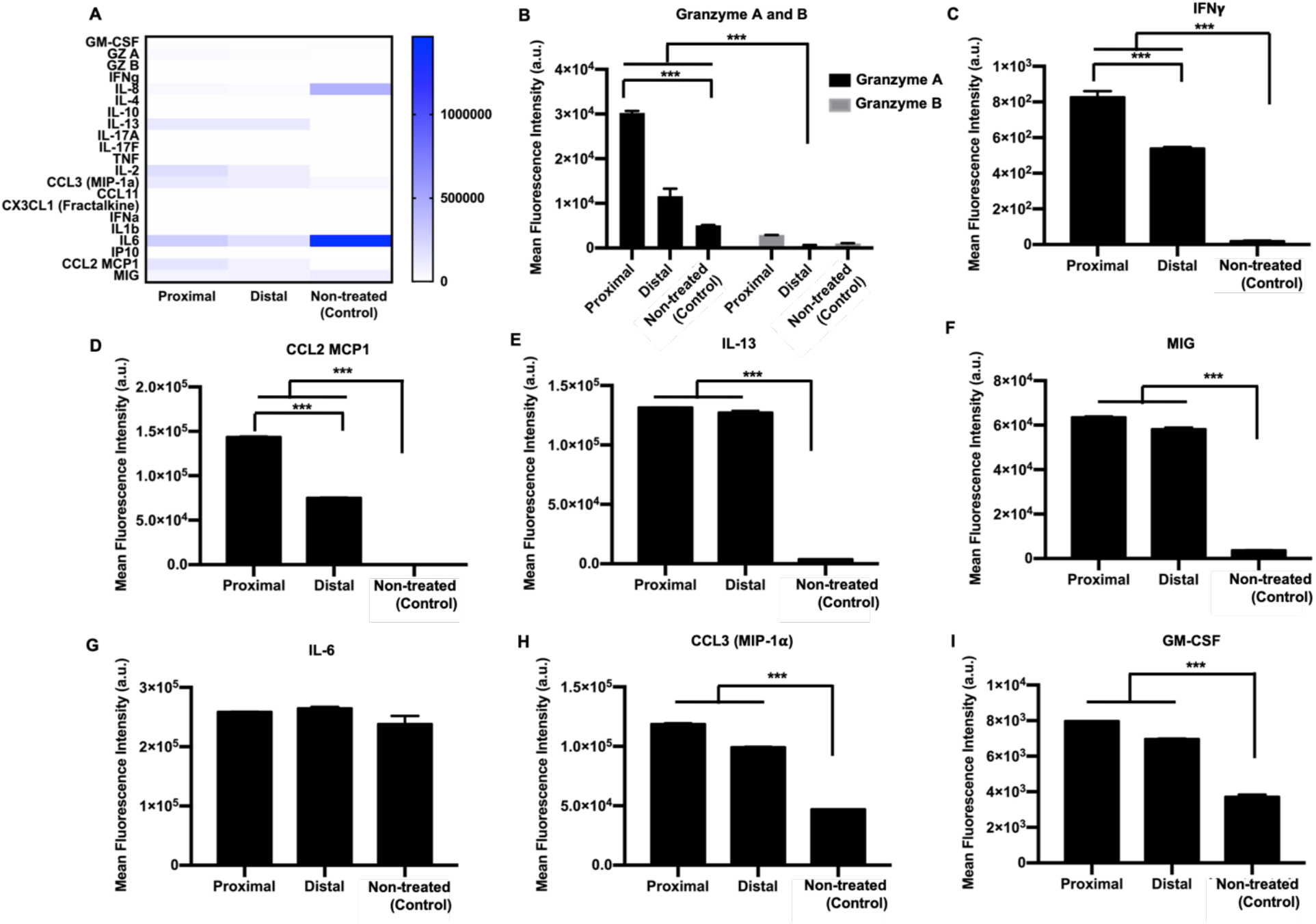
Supernatant analysis of 3D bioprinted tumor model employing MDA-MB-231/HDF heterotypic spheroids. (A) Heatmap of all cytokines and chemokines released in the supernatant post T cell treatment. Graphical representation of the mean fluorescence intensity of cytokines and chemokines secreted including (B) Granzyme A and B, (C) IFNγ, (D) CCL2, (E) IL-13, (F) MIG, (G) IL-6, (H) CCL3 and (I) GM-CSF (*w*=3, p*<0.05, p**<0.01, p***<0.001).

## DISCUSSIONS

In vitro tissue models that can efficiently recapitulate the dynamic immune-cancer microenvironment are greatly needed to understand the underlying tumor-immune interactions and to develop better immunotherapy approaches. In this study, 3D homotypic MDA-MB-231-only and heterotypic MDA-MB-231/HDF breast tumor models were developed to assess the efficacy of engineered T cells on tumor killing. CD8^+^ T cells engineered with MAIT TCRs were effectively able to lyse MDA-MB-231 cells in a MR1-dependent manner, over 3 days of culture. First, both the free-standing homotypic and heterotypic 3D tumor models exhibited a dose dependent increase in cancer cell death, indicating a greater eMAIT TCR-MR1 interaction. Extent of cell death was higher for free-standing homotypic model as compared to heterotypic model, which was indicative of higher immunoresistance in heterotypic cultures. Superior structural integrity of fibroblast containing tumors probably resulted in reduced immune cell infiltration. In accordance to the trend in cancer cell death, both free-standing homotypic and heterotypic models also exhibited a dose dependent increase in secretion of granzymes A and B, cytokines such as interferon gamma (IFNγ) and interleukin-13 (IL-13) and chemokine CCL3. Granzymes are cytotoxic molecules that are known to mediate cellular apoptosis by diffusing through perforin pores on the plasma membrane of target cells [19], [20]. IFNγ is also associated with antiproliferative, pro-apoptotic and antitumor mechanisms [21]. Thus, upregulation of both cytotoxic granzymes and IFNγ in homotypic and heterotypic tumor models substantiated T cell stimulation and induction of apoptosis in cancer cells. Secretion of proinflammatory cytokines IL-6 and IL-8 after eMAIT TCR-MR1 interaction, indicated potent immune response in both homotypic and heterotypic tumor models. Both IL-6 and IL-8 impact major hallmarks of cancer, including cancer cell proliferation, angiogenesis, metastasis, invasion as well as metabolism [22]–[24]. As IL-6 and IL-8 are not secreted by T cells, they were secreted by only MDA-MB-231 cells in the homotypic model where the secretion level of these factors decreased with increasing T cell treatment densities. In contrast, T cell treatment density exhibited no effect on IL-6 and IL-8 secretion of MDA-MB-231/HDF heterotypic tumors, indicating probable secretion of these immunosuppresive cytokines by HDFs [25], [26]. Cytokine granulocyte-macrophage colony-stimulating factor (GM-CSF) was expressed by both treated as well as non-treated tumors. It is often overexpressed in multiple tumor types but can also be expressed by T cells [23]. The homotypic tumor model exhibited a dose dependent increase in GM-CSF expression while T cell treatment density induced no differential effect in GM-CSF expression in the heterotypic tumor model, suggesting combined secretion of GM-CSF by both cancer and stromal cells [27].

Chemokines in the tumor microenvironment direct the movement of immune cells towards a tumor via autocrine or paracrine mechanisms [28]. Chemokine macrophage inflammatory protein-1 alpha (MIP-1α, CCL3) and monocyte chemoattractant protein-1 (CCL2/MCP-1) both play a pivotal role in immune cell stimulation and immune cell recruitment to intra-tumoral sites [29], [30]. CCL3 expression increased as the T cell dosage increased, for both homotypic and heterotypic tumor models. However, CCL2 was expressed only in the heterotypic tumor model suggesting a probable cross-talk between cancer cells and fibroblasts resulting in fibroblast activation and secretion of CCL2 [31]. Overexpression of CCL2 is known to promote tumor progression and provide immune resistance, as well as it could initiate infiltration of cytotoxic lymphocytes [32]. Additionally, monokine induced by gamma (MIG/CXCL9) is another chemokine whose expression decreased with increasing T cell treatment density in the homotypic tumor model in contrast to the heterotypic tumor model. MIG has been studied as a potential biomarker for breast tumor due to its abundance in breast cancer tissues as compared to normal tissues and correlated to enhanced immune cell infiltration of tumor cells [33]. Overexpression of MIG has shown to reduce tumor growth and metastasis [34]. As activated T cells could also induce MIG secretion, expression levels of MIG could be suggestive of an immune response post immunotherapy [35].

The scaffold-based homotypic tumor models also highlighted the efficiency of eMAIT TCR-MR1 interaction in eliminating a tumor target by freely migrating through the fibrin network. Subsequent upregulation of granzymes A and B, in conjuction with IFNγ, IL-13 and CCL-3 indicated T cell activation in these fibrin-based tumor models. Additionally, proinflammotory IL-6 and IL-8 cytokines were also enhanced post T cell treatment, thus generating a potent immune response. Additionally, secretion of GM-CSF and MIG probably further attracted the T cells to migrate towards the target MDA-MB-231 cells. Thus, the scaffold-based model emphasized the propensity of engineered T cells to maneuver a complex ECM-based microenvironment. Controlling the localization of T cells with respect to a MDA-MB-231/HDF tumor in the 3D bioprinted tumor model further revealed the importance of proximity in immune-cancer interactions. T cells bioprinted proximal (~250 μm) to the tumor inhibited cancer invasion more than the distal cultures (~650 μm). Proximal cultures presented higher expression of granzymes, IFNγ and CCL-3, all of which indicated higher immune cell activation as compared to distal cultures. Additionally, T cells bioprinted at a higher proximity to a heterotypic tumor exhibited a higher CCL2 production than the T cells bioprinted distally to the tumor. This suggested a probable role of collagen in mediating cancer cell-ECM interaction as well as immune activation resulting in subsequent cytokine secretion. These secreted cytokines can potentially be a post-treatment biomarker.

Studying immune-cancer interactions in different 3D tumor models with increasing complexity is essential for understanding the range of immune response and how it varies on introducing various complexities in 3D models [36], [37]. The tumor microenvironment is known to be immunosuppressive, with non-malignant cells aiding in tumor growth and progression[38]. Thus, it is important to incorporate different cell types, such as stromal cells and endothelial cells, in tumor models for a better understanding of the underlying mechanisms, which contribute to immune evasion and reduced anti-tumor response post immunotherapy. Our free-standing heterotypic tumor model as well the 3D bioprinted tumor model incorporates stromal cells along with cancer and immune cells. Importantly, difference in secretion levels of cytokines in different 3D models highlights the importance of studying immune-cancer interaction in the presence of stromal cells and ECM-based proteins, such as collagen and fibrin. Furthermore, with the 3D bioprinted model, we observed a difference in secretion of IFNγ, CCL2 and granzymes as the localization of T cells varied with respect to the tumor. This suggests paracrine mode of communication between T cells and cancer cells, as there was decreased secretion of these factors with increased distance between T cells and the tumor. Although this presented study explored 3D tumor models for TCR therapy, which requires MHC molecules for T cell-cancer cell recognition, more physiologically-relevant tumor models need to be established to explore other immune-modulatory therapies, such as chimeric antigen receptor (CAR) therapies. CAR therapies directly recognize tumor antigens and 3D tumor models with increasing complexity could enable deeper understanding of the anti tumor efficacy of CARs and thereby accelerate the development of approaches to improve cancer immunotherapy.

## CONCLUSION

In this study, eMAIT TCR-MR1 interaction was explored employing different 3D breast tumor models to understand the efficiency of TCR-modified engineered T cells in eradicating tumors. Engineered T cells effectively eliminated cancer cells in 3 days of culture in 5-ARU as well as MR1 dependent manner. They were activated when cultured with MR1 expressing MDA-MB-231 cells and activated T cells secreted a range of cytokines, whose expression levels varied between homotypic and heterotypic spheroids. Non-engineered conventional CD8^+^ T cells did not affect the viability of MDA-MB-231 cells nor did the inactive 6-FP compound. Furthermore, bioprinting of T cells and tumor spheroids in a localized manner, exhibited that T cell activation was affected by the distance between T cells and cancer cells suggesting paracrine mechanism of communication. Overall, the measured immune response from these 3D models in response to immunotherapy, shed light on the importance of different tumor models in studying underlying immune-cancer interactions. Our study also substantiates the importance of studying eMAIT TCR-MR1 interactions as a potential immunotherapy modality since numerous tumors express the MR1 protein.

## ACKNOWLEDGEMENT

We are grateful to late Dr. N. Zavazava (Department of Internal Medicine at the University of Iowa, Iowa city, IA) for providing HDFs and Dr. Danny Welch (University of Kansas, Kansas City, KA) for providing MDA-MB-231 cells. We acknowledge Bugra Ayan, a former graduate student at Penn State, for his assistance in designing the CAD model and 3D printing PLA frames. We also acknowledge the support from The Huck Institutes of Life Sciences and Materials Research Institute for providing facilities for characterization of experiments. This work was supported by NSF awards 1914885 (I.T.O.) and 1624515 (I.T.O.) and NIH award R21 CA224422 01A1 (I.T.O. and D.U.)

## AUTHOR CONTRIBUTIONS

M.D., D.U. and I.T.O developed the ideas and designed the experimental plan. M.D., M.H.K., M.N., E.K., L.K. and M.D. performed the experiments. M.D. took the lead in writing the manuscript. All authors provided critical feedback and helped shape the research, analysis and manuscript, and approved the content of the manuscript.

## DATA AVAILIBILITY

All data needed to evaluate the conclusions in the paper are present in the paper. Additional data related to this paper may be requested from the authors.

