## Supplementary Information for "Development of 3D breast cancer models with human T cells expressing engineered MAIT cell receptors"

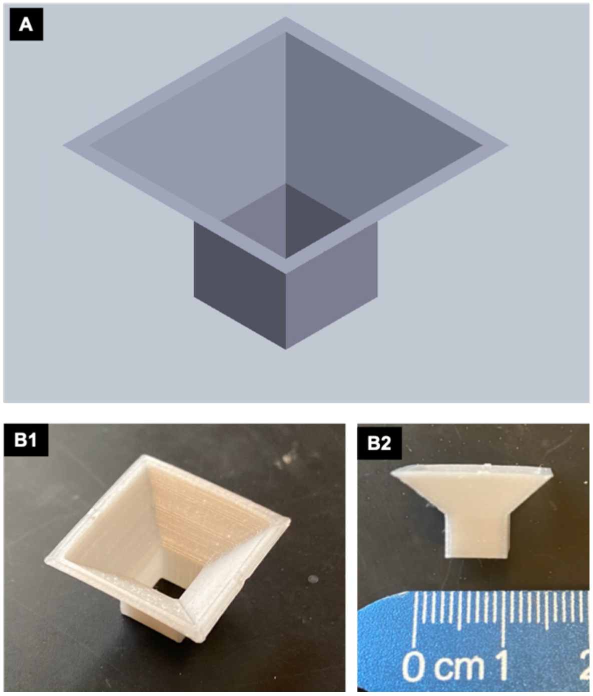


**Fig S1**: (A) Computer aided design (CAD) model of the 3D box. (B1-B2) PLA-based 3D printed box frame.

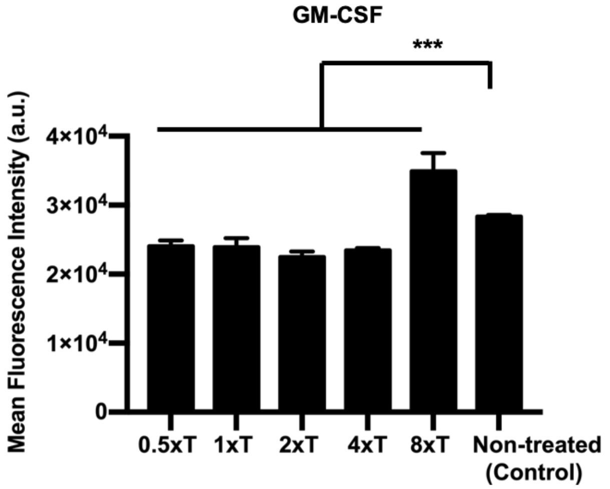


**Fig. S2**. Graphical representation of the mean fluorescence intensity of GM-CSF secretion in heterotypic MDA-MB-231/HDF spheroids after the T cell treatment (*n*=3, p*<0.05, p**<0.01, p***<0.001).

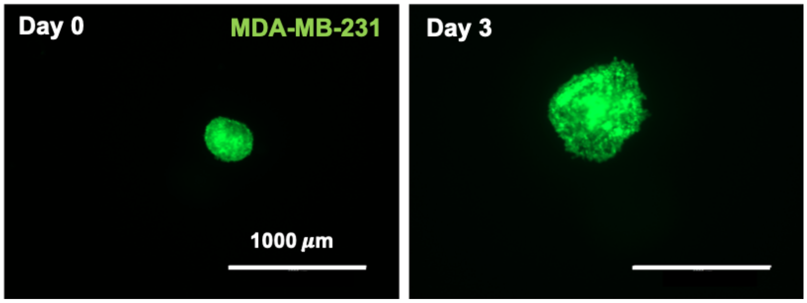


**Fig. S3**: Fluorescent image of 3D bioprinted tumor spheroids in collagen without employing T cell bioprinting, at Day 0 (right after bioprinting of the spheroid) and Day 3. This served as the non-treated control group.

**Supplementary Videos**

**Supplementary Video S1**: Time lapse video of T cells migrating into the fibrin network. The T cells were color-coded based on their 'Z' locations in the 3D model.

**Supplementary Video S2**: Cancer cells were encapsulated in fibrin and T cells were exposed from the top. This video highlights the fate of cancer cells during the T cell treatment. The cancer cells have been color-coded based on their 'Z' location. The 3D model was divided into three zones, namely, top, middle and bottom zone. Cells in the top zone exhibited reduced fluorescence on treatment with cytotoxic T cells whereas, cells in the bottom zone proliferated and increased in GFP intensity.

**Supplementary Video S3**: Time lapse video of MDA-MB-231 spheroid encapsulated in fibrin with engineered T cells. Tumor cells migrate and invade into its surrounding matrix. T cells were also able to migrate through the fibrin network and find its target.
